# Prostaglandin PGE2 receptor EP4 regulates microglial phagocytosis and increases susceptibility to diet-induced obesity

**DOI:** 10.1101/2021.11.18.469049

**Authors:** Anzela Niraula, Rachael D. Fasnacht, Kelly M. Ness, Jeremy M. Frey, Mauricio D. Dorfman, Joshua P. Thaler

## Abstract

**Background:** In rodents, susceptibility to diet-induced obesity requires microglial activation, but the molecular components of this pathway remain incompletely defined. Prostaglandin E2 (PGE2) levels increase in the mediobasal hypothalamus during high fat diet (HFD) feeding, and the PGE2 receptor EP4 regulates microglial activation state and phagocytic activity, suggesting a potential role for microglial EP4 signaling in obesity pathogenesis.

**Method:** Metabolic phenotyping, as assessed by body weight, energy expenditure, glucose, and insulin tolerance, was performed in microglia-specific EP4 knockout (MG-EP4 KO) mice and littermate controls on HFD. Morphological and gene expression analysis of microglia, and a histological survey of microglia-neuron interactions in the arcuate nucleus was performed. Phagocytosis was assessed using *in vivo* and *in vitro* pharmacological techniques.

**Results:** Microglial EP4 deletion markedly reduced weight gain and food intake in response to HFD feeding. In correspondence with this lean phenotype, insulin sensitivity was also improved in the HFD-fed MG-EP4 KO mice though glucose tolerance remained surprisingly unaffected. Mechanistically, EP4-deficient microglia showed an attenuated phagocytic state marked by reduced CD68 expression and fewer contacts with POMC neuron soma and processes. These cellular changes observed in the microglial EP4 knockout mice corresponded with an increased density of POMC neurites extending into the paraventricular nucleus.

**Conclusion:** These findings reveal that microglial EP4 signaling promotes body weight gain and insulin resistance during HFD feeding. Furthermore, the data suggest that curbing microglial phagocytic function may preserve POMC cytoarchitecture and PVN input to limit overconsumption during diet-induced obesity.

## Introduction

Hypothalamic regulation of energy balance maintains systemic metabolism and body weight homeostasis. A key neurocircuit in the control of food intake and energy balance is the melanocortin system, comprised of proopiomelanocortin (POMC) neurons and agouti-related peptide (AgRP) neurons in the arcuate nucleus. These mutually inhibitory neurons integrate hormonal and nutritional signals of energy status to calibrate the activity of downstream melanocortin-receptor-expressing neurons in the paraventricular nucleus (PVN), thereby controlling sympathetic tone and consummatory drive [1]. POMC neurons, in particular, operate over long time frames to limit weight gain by reducing food intake and increasing energy expenditure, and dysfunction in this neuronal subset is thought to be a critical component of obesity pathogenesis [2].

High fat diet (HFD) feeding triggers an inflammatory response marked by gliosis in the arcuate nucleus. Under homeostatic conditions, microglia exhibit a ramified morphology with long processes constantly surveilling the brain microenvironment. However, following perturbations such as HFD, these cells undergo rapid morphological changes, marked by a more ameboid-shaped cell body with thicker processes [3]. Importantly, this structural remodeling is indicative of cellular activation, frequently characterized by production of cytokines and immune mediators, such as interleukin-1 beta (IL1β) and tumor necrosis factor alpha (TNFα), as well as increased phagocytic activity [3–6]. These microglial changes, collectively referred to as gliosis, contribute to susceptibility to diet-induced obesity (DIO) because pharmacological depletion of microglia or genetic disruption of their NF-κB (nuclear factor kappa-light-chain-enhancer of activated B cells) signaling pathway reduces food intake and weight gain during HFD feeding [3, 6]. However, the downstream mechanisms triggered by HFD feeding to alter microglial activity remain poorly characterized.

Prostaglandins are signaling lipids derived from arachidonic acid that are present throughout the body and perform an array of functions in health and disease [7]. Prostaglandin E2 (PGE2) is a key inflammatory mediator that regulates apoptosis, angiogenesis, and cell proliferation, but is also implicated in diseases such as stroke, neurodegenerative conditions, and cancer [7, 8]. PGE2 signals through four G protein-coupled receptors (EP1-4), the downstream effects of which vary across tissues and conditions [8]. While their role in DIO is unclear, PGE2 levels are elevated in the arcuate nucleus of HFD-fed mice [9], which could modulate microglial activity through the EP receptors. EP4 is highly expressed in microglia and generally serves to constrain the microglial response to inflammatory stimuli such as lipopolysaccharide (LPS) and beta amyloid plaque in rodent models of Alzheimer’s Disease [10, 11]. However, the effect of EP4 signaling can vary based on context and timing. For example, in experimental autoimmune encephalomyelitis (EAE) – a rodent model of multiple sclerosis – EP4 signaling promotes the development of pathology in early stages of disease but reduces severity at later time points [12]. Since EP4-null mice develop a lipodystrophic phenotype with a shortened lifespan [13], it is critical to study EP4 function using cell-specific methods to determine its precise role in metabolic homeostasis.

In the current study, we delineated the role of microglial EP4 signaling in energy balance and glucose homeostasis. Microglial EP4 deletion markedly reduced weight gain during HFD feeding as a result of lower energy intake with no effect on chow-fed mice. Notably, EP4-deficient microglia showed evidence of less microglial phagocytic activity, which was associated with fewer microglia-neuron cell-cell contacts and greater POMC neuron fiber density in the PVN, suggesting that the lean phenotype in the microglial EP4 knockout mice results from preservation of the melanocortin system during DIO.

## Methods

### Mice

Microglia-specific EP4 knockout (MG-EP4 KO) mice were generated by crossing the CX3CR1^(CreERT2/+)^ (B6.129P2(Cg)-Cx3cr1^tm2.1(cre/ERT2)Litt^/WganJ; Jackson stock no. 021160) line with the Ptger4^fl/fl^ line (originally generated by Matthew and Richard Breyer, and generously provided by Karin Bornfeldt, University of Washington). Along with the Cre allele, the CX3CR1^(CreERT2/+)^ mice contain an insert for enhanced yellow fluorescent protein (EYFP), thus allowing for fluorescence-mediated tracking of Cre in CX3CR1-expressing cells. The resulting CX3CR1^CreERT2(+/−)^::Ptger4^fl/fl^ offspring were injected at 6 weeks of age with 2 doses of tamoxifen (T5648, Sigma, 4 mg s.c., prepared in corn oil) 48 hours apart. Following a 4-week period to allow replenishment of peripheral myeloid cells [14], mice were single housed, and placed on a 60% kcal HFD (D12492i, Research Diets). CX3CR1^CreERT2(−/−)^::Ptger4^fl/fl^ littermates were used as wildtype controls and subjected to the same tamoxifen regimen. Chow-fed mice were placed on the standard diet (5053, LabDiet). All mice had *ad libitum* access to food and water except during fasting experiments, and were housed in a temperature-controlled room with 14h/10h light/dark cycle. All experiments and procedures were performed in accordance with NIH Guidelines for Care and Use of Animals, and were approved by the Institutional Animal Care and Use Committee at the University of Washington.

### Fluorescent Activated Cell Sorting (FACS)

At 3 weeks on HFD, mice were anesthetized and transcardially perfused with cold PBS. Brains were collected and homogenized on ice using a Dounce homogenizer. The centrifuged pellet was resuspended in 33% Stock Isotonic Percoll (GE Healthcare), and the myelin-free pellet was recovered following centrifugation. Following resuspension in PBS and passage through a 70μm strainer, cells were incubated in Fc block solution (Bio Rad, Cat# BUF041A) and PerCP-conjugated anti-CD11b antibody (BD Pharminogen, Cat# 550993). After PBS washes, cells were stained with DAPI and sorted on the FACS Aria II sorter (Becton Dickinson) at the University of Washington Cell Analysis Facility. CD11b+ cells were collected in RNA lysis buffer (Qiagen) and processed for real time quantitative PCR analysis.

### Magnetic Activated Cell Sorting (MACS)

Brains from chow-fed mice were isolated and processed in a manner similar to FACS. Following filtration through the strainer, the cell suspension was incubated with anti-CD11b MicroBeads (130-049-601, Miltenyi Biotec) at 4°C for 15 minutes. The cells were passed through a column using a MiniMACS™ Separator, and the CD11b- and CD11b+ fractions were collected in separate tubes. Cells were stained with DAPI, placed on glass slides, and coverslipped. The EYFP signal was visualized using a Nikon epifluorescence microscope (Eclipse E600), and images captured with a mounted Nikon camera.

### Body composition analysis

Body composition analysis, including fat mass and lean mass measurements, were performed using EchoMRI immediately prior to indirect calorimetry in awake, conscious mice at the University of Washington, Nutrition Obesity Research Center (NORC) Energy Balance Core.

### Indirect Calorimetry

Indirect calorimetry was performed at the University of Washington NORC facility. Mice were acclimated to metabolic cages connected to an indirect calorimetry system (Sable Systems, Promethion), and monitored for energy expenditure over multiple light and dark cycles. Data were collected continuously at 5 minute intervals, and measures of O_2_ consumption, CO_2_ production, ambulatory activity, and food and water intake were recorded. Respiratory quotient (RQ) was calculated as a ratio of CO_2_ production to O_2_ consumption. Acquired data were binned into hourly datapoints for analysis and presentation. Energy expenditure data for HFD mice was adjusted for body weight differences using ANCOVA.

### Intraperitoneal Glucose Tolerance Test (IPGTT)

Following a 4-hour food withdrawal period, mice were injected with a bolus of glucose (2 g/kg i.p., 30% dextrose prepared in saline). Glucose values were obtained via tail vein bleeding, and measurements performed at the indicated timepoints using a glucometer (Freestyle Freedom Lite).

### Intraperitoneal Insulin Tolerance Test (IPITT)

Following a 4-hour food withdrawal period, mice were injected with insulin (0.9 U/kg ip prepared in saline, Humulin). Glucose values were obtained via tail vein bleeding, and measurements performed at the indicated timepoints using a glucometer (Freestyle Freedom Lite). The glucose measurements were normalized to the baseline (time 0) value for each mouse.

### Insulin ELISA

Plasma samples were collected following a 16 hour overnight fast. Insulin ELISA was performed following manufacturer’s instructions (Crystal Chem, Cat# 90080).

### Leptin ELISA

Plasma samples were collected at baseline (non-fasting), and ELISA was performed following manufacturer’s instructions (Crystal Chem, Cat# 90030).

### Intracerebroventricular Administration of EP4 Agonist

Mice were placed under 3% isoflurane to induce sedation, and maintained on 1.5% isoflurane during surgical implantation of steel guide cannulas (Alzet, DURECT Corp.) into the lateral ventricle (coordinates x=+1.3mm, y=−0.7mm, z=−1.3mm), and allowed to recover for 2 weeks prior to experiments. During intracerebroventricular (i.c.v.) infusion, a 1 μL solution of the EP4 agonist, L-902,688, (25 nmol; Cayman Chemicals) was administered using an injector connected via tubing to a 10 μL Hamilton syringe. Vehicle mice received the corresponding vehicle solution (50% DMSO prepared in sterile saline). Mice received two i.c.v. infusions (72 hours apart) of vehicle or drug. Thirty minutes after the last infusion, mice were perfused, and brains collected for immunofluorescent labeling as described below.

### Immunofluorescent Labeling and Microscopy

Mice were anesthetized with intraperitoneal injections of ketamine (100 mg/kg) and xylazine (10 mg/kg), and transcardially perfused with ice cold PBS followed by 4% paraformaldehyde. Brains were collected and fixed overnight in 4% paraformaldehyde, then transferred to 25% sucrose for 48 hours, and frozen until processing. A freezing stage microtome (Leica SM2010R) was used to section the brains, and 25 μm free-floating sections were collected for immunolabeling. Sections were blocked using 5% normal donkey serum (Jackson Immunoresearch), followed by overnight incubation with primary antibodies: rabbit anti-Iba1 (Wako, 1:1000), goat anti-Iba1 (Abcam, 1:1000), rat anti-CD68 (BioRad, 1:1000), and rabbit anti-β-endorphin (Phoenix Pharmaceuticals, 1:5000). Washes were performed followed by subsequent incubations with secondary antibodies: donkey anti-rabbit Alexa Fluor 488 (ThermoFisher, 1:500), donkey anti-rat Alexa Fluor 594 (ThermoFisher, 1:500), donkey anti-rabbit Alexa Fluor 594 (ThermoFisher, 1:500), and donkey anti-rabbit Alexa Fluor 647 (ThermoFisher, 1:500). DAPI (1:10000) was used for nuclear staining. Sections were mounted on slides and coverslipped in the presence of mounting medium (Fluoromount-G, ThermoFisher).

Samples were visualized using a Keyence BZ-X800 inverted microscope. CD68 and Iba1 co-labeling images were captured at 20x magnification. Images for microglial morphology (Fig. 3A) and microglia-neuron apposition analysis (Fig. 4A) were captured at 60x using the sectioning feature to generate a z stack (1 μm slice; 30 slices per stack) and processed using the maximum projection intensity feature.

### Microglial Sholl and Morphology Analysis

Sholl analysis was performed using the Sholl plugin on Fiji (NIH ImageJ FIJI). Briefly, images were transformed into binary and thresholded to clearly visualize microglia and processes. 2 microglia per image (field of view) in 2 images per mouse were analyzed. Each cropped microglia was reconstructed via tracing using the Simple Neurite Tracer plugin on FIJI. A fixed ROI was drawn at the soma of the reconstructed microglia. The start radius was set at 0.05 arbitrary units (a.u.) and end radius 4 a.u. with step size of 0.2. Number of intersections were automatically generated by the software. Data were averaged across microglia, and an area under the curve was calculated for each mouse. Soma size, circularity, and Feret length were calculated using the ROI feature on FIJI.

### In Vitro Phagocytosis Assay

BV2 cells (kind gift from Dr. Suman Jayadev, University of Washington) were cultured and passed three times prior to experiment. Cells were plated onto a 12 well-plate (50,000 cells per well) and incubated in DMEM supplemented with 5% fetal bovine serum and 1% pen/strep at 37°C/5% CO2 overnight. Upon reaching confluency, cells were serum-starved 24 hours prior to the experiment. On the day of the experiment, cells were treated with varying concentrations of the EP4 agonist L-902,688 (Cayman Chemical) for an hour, followed by incubation with fluorescent-tagged carboxylate-coated 2 um size latex beads (L3030, Sigma) for 3 hours. Cells were detached from the wells with light trypsinization, followed by washes, and flow analysis (Becton Dickinson, Canto). DAPI was used to exclude dead cells. Cells positive for phycoerythrin were gated and quantified as a percentage of total viable cells.

### Statistics

Statistical analyses were performed on GraphPad Prism 9.0.2 using unpaired two-tailed Student’s t tests for group-wise comparisons. Body weight gain and cumulative food intake data were analyzed using a two-way analysis of variance (ANOVA), and post hoc tests were performed using Šidák test to correct for multiple comparisons. Indirect calorimetry data for heat production was analyzed using analysis of co-variance (ANCOVA) with body mass as covariate. Results are presented as mean ± SEM. P values <.05 were considered statistically significant.

## Results

### Microglial EP4 deletion reduced susceptibility to DIO

A recent study reported an increase in PGE2 levels in the arcuate nucleus following high fat diet (HFD) feeding [9]. Interestingly, we found a near-significant increase in expression of the receptor EP4 (encoded by the gene, *Ptger4*) in isolated microglia but not whole hypothalamus from HFD-fed mice (Supplementary Fig. S1A), suggesting a potential role for EP4 in dietary modulation of microglial function. To explore this possibility further, we generated CX3CR1^CreERT2(+/−)^ ^IRES^ ^EYFP^::Ptger4^fl/fl^ mice (MG-EP4 KO) and CX3CR1^CreERT2(−/−)^::Ptger4^fl/fl^ littermate controls (WT), which are predicted to develop EP4 deficiency in microglia upon tamoxifen treatment. As expected, tamoxifen treatment led to a significant reduction (~79%) in microglial *Ptger4* expression in the MG-EP4 KO mice compared with WT littermates (Supplementary Fig. S1B). To confirm the cellular specificity of the EP4 knockout, we isolated CD11b+ cells (CNS microglia/myeloid) and CD11b-cells (neurons and other glia) from WT and MG-EP4 KO mice using Magnetic Activated Cell Sorting (MACS). Given that *EYFP* and *CreERT2* are co-transcribed from a bicistronic mRNA cassette knocked into the *Cx3cr1* locus, EYFP fluorescence serves as a faithful marker of *Cre* expression. We found EYFP only in CD11b+ cells and not in any population of CD11b-cells (Supplementary Fig. S1C), indicating that EP4 deletion occurred exclusively in the CNS myeloid compartment of the MG-EP4 KO mice. Mice were born at a Mendelian ratio (data not shown) and had no difference in body weight for at least 90 days on standard chow (Supplementary Fig. S1D), suggesting that microglial EP4 signaling is dispensable for energy balance under normal physiologic conditions.

Given the rapid activation of microglia during HFD feeding, we hypothesized that microglial EP4 deletion would affect the susceptibility to DIO via alterations to microglial functions. Indeed, MG-EP4 KO mice had a marked protection from HFD-induced weight gain (Fig. 1A) and fat mass gain (Fig. 1B) compared to the WT mice. Though lean mass was modestly reduced as well (Fig. 1B), lower plasma leptin levels in KO mice (Fig. 1C) indicated a strong effect of microglial EP4 deletion on the regulation of adiposity. Tracking of food intake over the course of 105 days of HFD feeding revealed significantly lower cumulative food intake in the MG-EP4 KO animals (Fig. 1D). Examining food intake in metabolic cages during calorimetry revealed that the MG-EP4 KO mice consumed less HFD in both light and dark cycles (Fig. 1E-G), resulting in reduced total daily intake (Fig. 1G). Surprisingly, body weight-normalized energy expenditure was also lower in MG-EP4 KO mice (Fig. 1H-J), suggesting a compensatory metabolic mechanism to offset hypophagia. However, with respiratory quotient closer to 0.7 (the level indicative of fat utilization) in KO mice (Fig. 1K and L), this adaptation is apparently inadequate, resulting in consistent body weight differences between genotypes. Overall, these results indicate that microglial EP4 signaling promotes HFD-induced hyperphagia and susceptibility to diet-induced obesity.

**Figure 1:**
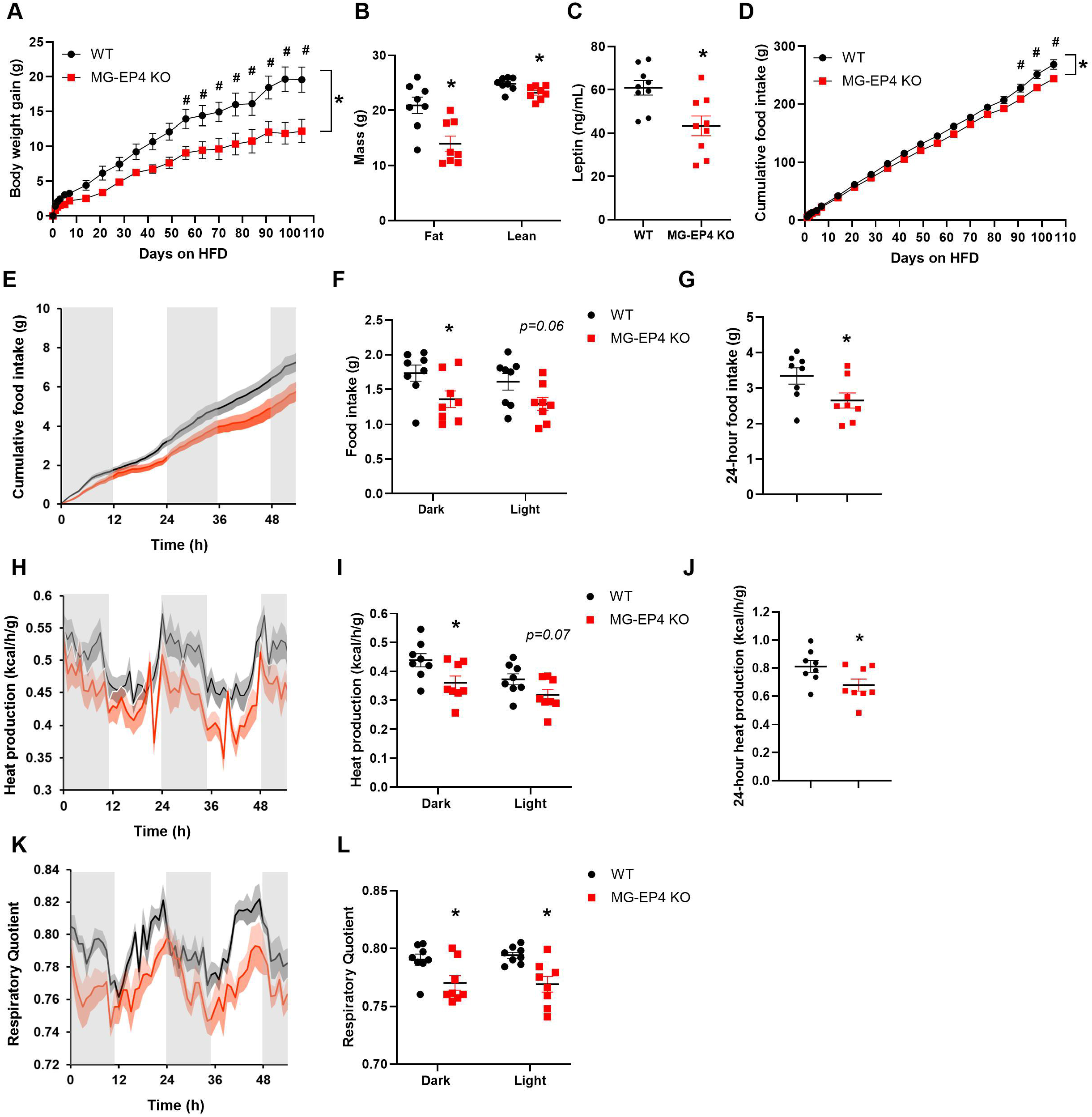
Microglial EP4 deletion reduced susceptibility to DIO. (A) Body weight gain, (B) fat mass and lean mass, (C) plasma leptin, and (D) cumulative food intake in male WT and MG-EP4 KO mice on HFD for 16 weeks (n=8-9). Continuous measurements of (E) cumulative food intake, (F) binned dark cycle and light cycle food intake, and (G) total 24h food intake. (H) Heat production traces, (I) quantification of dark cycle and light cycle, and (J) total heat production. (K) Respiratory quotient traces and (L) quantification of dark cycle and light cycle respiratory quotient following 16 weeks on HFD. (n=8). Body weight gain and cumulative food intake data analyzed by 2-way ANOVA with Šidák test. *p < .05 for genotype x time interaction, #p < .05 for post hoc analysis. Data in (H-J) analyzed by ANCOVA with body mass as covariate. Values represent mean ±SEM. *p < .05, Student’s *t test.*

### Microglial EP4 deletion improved insulin tolerance on HFD

Given the lean phenotype induced by microglial EP4 deletion, we hypothesized that obesity-associated glucose intolerance would be improved in MG-EP4 KO mice. However, no differences in glucose tolerance between genotypes were observed at 12 weeks on HFD (Fig. 2A). Instead, MG-EP4 KO mice showed lower fasting plasma insulin (Fig. 2B) with no difference in fasting glucose, (Fig. 2C) indicating increased insulin sensitivity. Similar results were obtained at 22-23 week HFD with equivalent glucose tolerance (Fig. 2D) but improved insulin tolerance in MG-EP4 KO mice compared to WT controls (Fig. 2E). These findings suggest that microglial EP4 deletion improves insulin sensitivity but leaves glucose tolerance unaffected presumably from lower insulin production.

**Figure 2:**
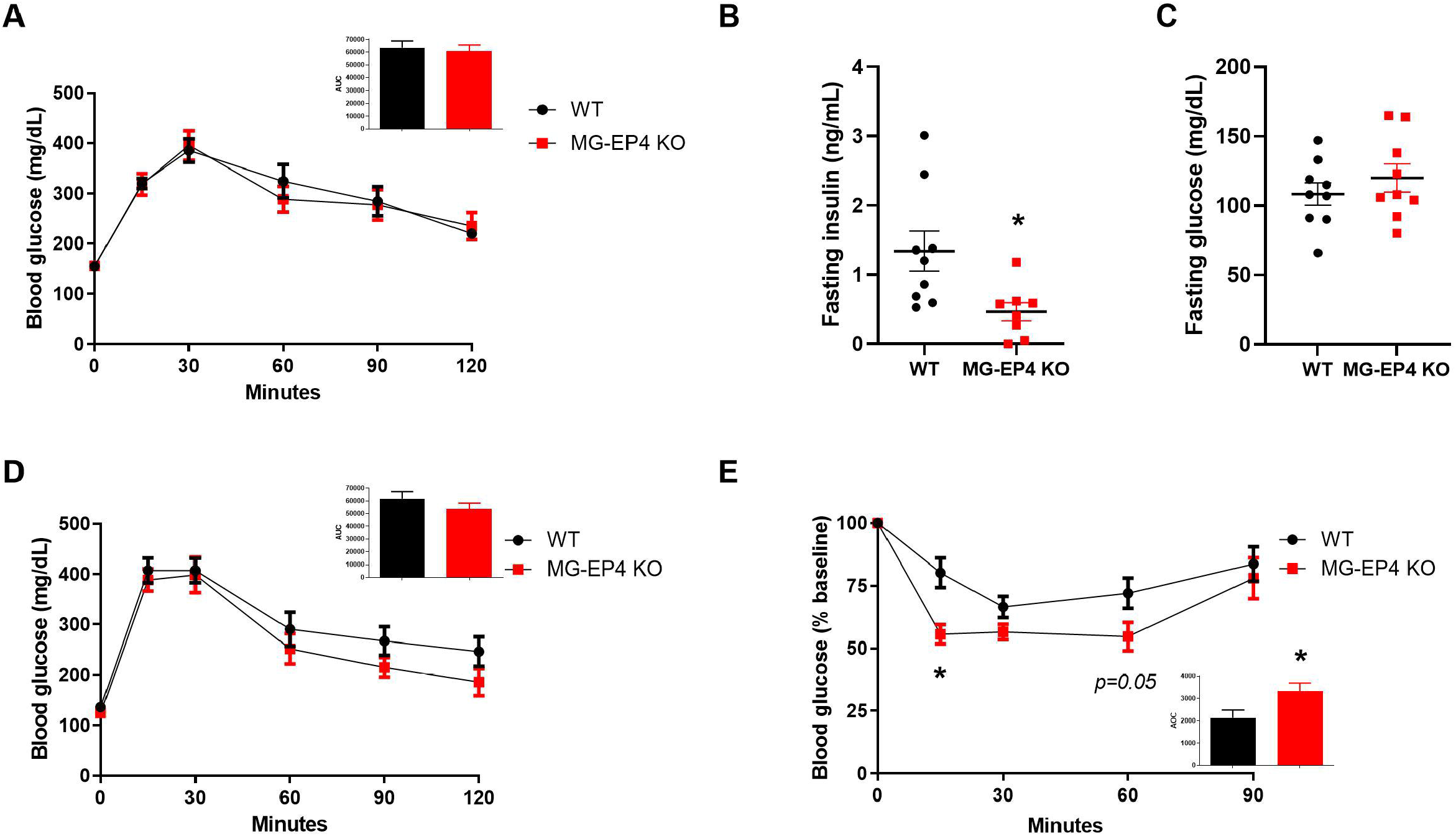
Microglial EP4 deletion improved insulin tolerance on HFD. (A) Intraperitoneal glucose tolerance test (2 g/kg i.p.) and area under curve (AUC) analysis (inset) from male WT and MG-EP4 KO mice following 12 weeks of HFD feeding (n=9). Fasting plasma insulin (B) and glucose (C) levels at 12 weeks on HFD. (D) Intraperitoneal glucose tolerance test with AUC (inset) at 22 weeks of HFD and (E) insulin tolerance test (0.9U/kg) with area over curve at 23 weeks of HFD (n=9). Values represent mean ±SEM. *p < .05, Student’s *t test.*

**Figure 3:**
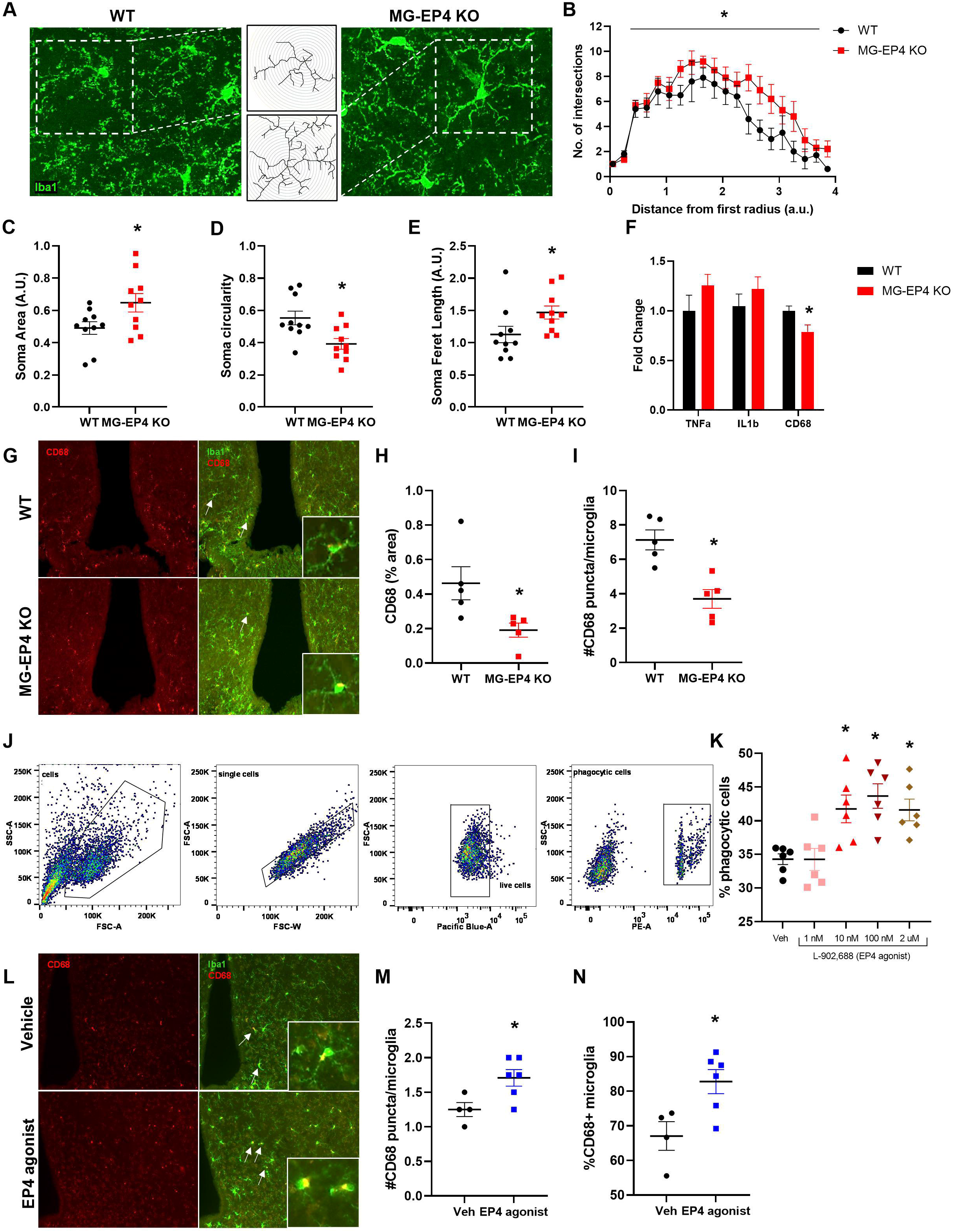
Microglial EP4 deletion reversed key HFD-induced alterations to microglial attributes. (A) Representative maximum intensity projection images of fluorescently labeled Iba1+ microglia from WT and MG-EP4 KO mice following 3 weeks of HFD. Insets represent reconstructed cells in the middle panel (top = WT, bottom = MG-EP4 KO). (B) Area under curve comparison between WT and MG-EP4 KO mice following Sholl analysis of microglial process intersections. (C) Cell soma area, (D) circularity, and (E) soma Feret length. Datapoints represent 2 sections/mouse; n=5). (F) Quantitative real time PCR measurements of genes from FAC-sorted microglia following 3 weeks on HFD (n=8-10). (G) Representative images of CD68 and Iba1 immunofluorescence labeling from WT and MG-EP4 KO brains following 5 weeks on HFD. Arrows represent CD68+ microglia from the mediobasal hypothalamus also depicted in the insets. Quantification of (H) CD68 percent area and (I) number of CD68+ puncta per microglia (n=5). (J) Representative flow cytometry plots of BV2 cells treated for 3 hours with 2 μm phycoerythrin-coated latex beads in presence of various concentrations of the EP4 agonist (L-902,688). (K) Bead-positive cells out of total live cells quantified as percentage phagocytic cells (n=3 from 2 technical replicates). (L) Representative images of CD68 and Iba1 immunofluorescence in the mediobasal hypothalamus following intracerebroventricular administration of the L-902,688 (25 nmol) or vehicle in chow-fed wildtype mice. Arrows represent CD68+ microglia also depicted in the insets. (M) Quantification of CD68+ puncta per microglia and (N) percentage of CD68+ microglia (n=4-6). Values represent mean ±SEM. *p < .05, Student’s *t test.*

**Figure 4:**
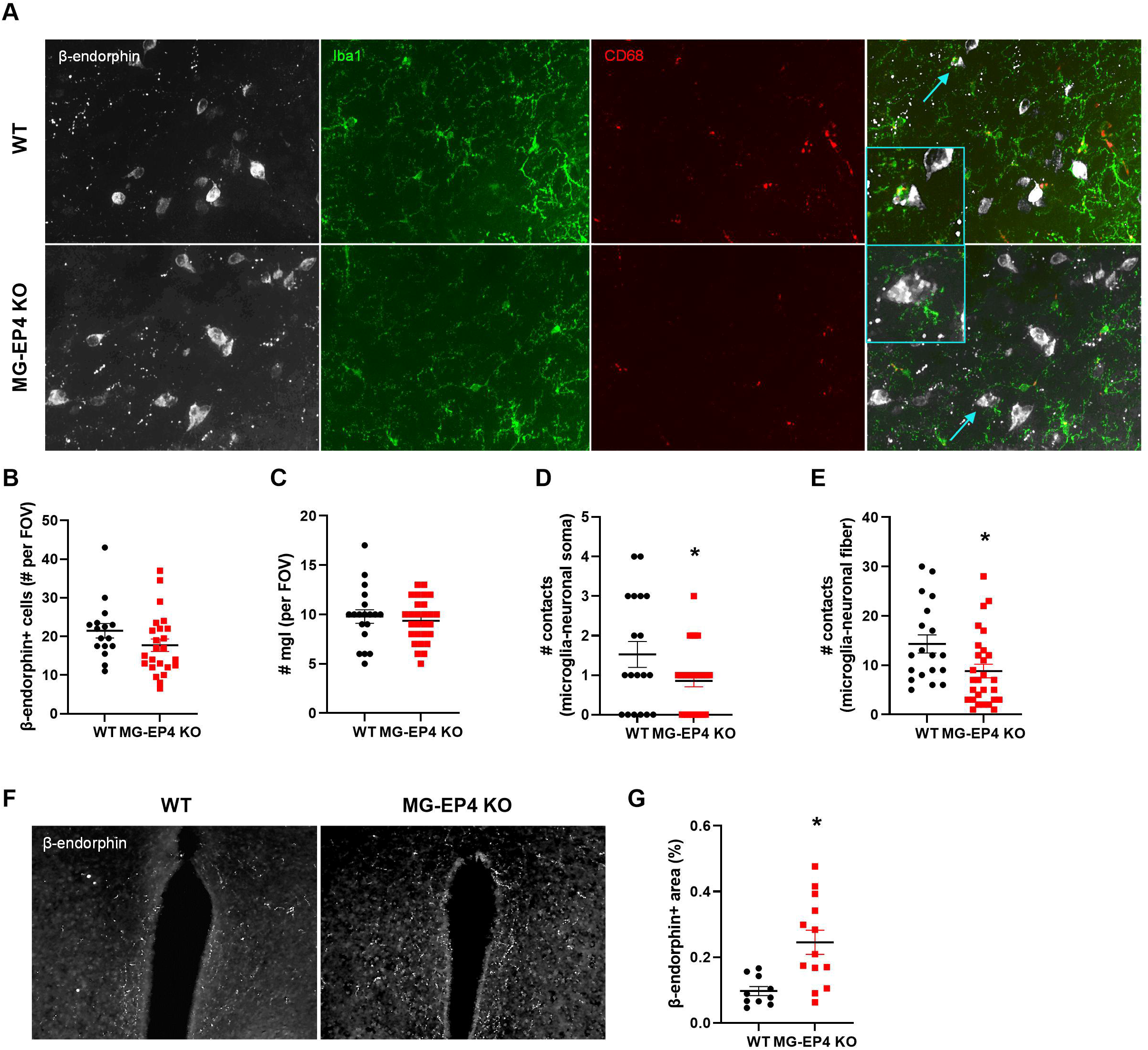
Microglial EP4 deficiency preserved POMC neuron projections to the PVN during HFD feeding. (A) Representative images of POMC neurons marked by β-endorphin positivity (grey), Iba1+ microglia (green), and CD68+ puncta (red) in the arcuate nucleus of the mice following 3 weeks on HFD. Blue arrows in the rightmost panels represent microglia-neuron appositions depicted in the insets. Quantification of (B) β-endorphin+ cell number and (C) microglia number per field of view (FOV), (D) number of microglia-neuronal soma contacts, and (E) microglia-neuronal process contacts in the arcuate nucleus. Datapoints represent 3 sections/mouse (n=5-7). (F) Representative images and (G) quantification of β-endorphin+ area in the paraventricular nucleus. Datapoints represent 2 sections/mouse (n=5-7). Values represent mean ±SEM. *p < .05, Student’s *t test.*

### Microglial EP4 deletion reversed key HFD-induced alterations to microglial attributes

Our findings demonstrated a positive association between microglial EP4 deletion and reduced susceptibility to weight gain and insulin resistance during HFD feeding. To begin to elucidate the mechanisms underlying this phenotype, we examined microglial properties that are altered during HFD exposure. Morphological assessment of microglia using Sholl analysis following 3 weeks of HFD revealed that MG-EP4 KO mice had more intersections of arcuate nucleus microglial processes with Sholl radii than WT controls, indicating a more ramified structure of EP4-deficient microglia (Fig. 3A&B). MG-EP4 KO microglia also had larger cell somas (Fig. 3C), lower circularity indices (Fig. 3D), and longer Feret lengths (i.e. distance between the two farthest points of the cell soma) (Fig. 3E). Taken together, these cytological features suggest that EP4 deficiency renders microglia less susceptible to HFD-mediated activation.

Based on the morphological analysis above, we predicted that MG-EP4 KO microglia would have reduced inflammatory gene expression consistent with a less activated profile. Unexpectedly, however, microglial *Il1*β and *Tnf*α mRNA levels did not differ between genotypes after 3 weeks of HFD feeding (Fig. 3F). Instead, expression of the *Cd68,* a marker of phagocytosis, was significantly reduced in the MG-EP4 KO microglia compared to the WT (Fig. 3F). Examining tissue sections of the mediobasal hypothalamus, we observed that CD68 immunoreactivity was diminished in the arcuate nucleus in the MG-EP4 KO mice (% area; Fig. 3G and H). As a lysosomal membrane-associated glycoprotein [15], CD68 produces a punctate staining pattern under immunofluorescence (see Fig. 3G inset). Quantification of high magnification images revealed fewer CD68+ puncta localized in MG-EP4 KO microglia (Fig. 3I).

Reduced CD68 in KO microglia suggests that microglial phagocytosis is regulated by EP4 signaling. To test this hypothesis *in vitro*, we treated the murine microglial cell line BV2 with the selective EP4 agonist, L-902,688 and measured phagocytosis of fluorescently-labeled carboxylate-coated latex beads using flow cytometry. EP4 agonist treatment increased the percentage of phagocytic cells at concentrations above 1 nM (Fig. 3J and K). Similarly, central administration of L-902,688 (25 nmol, i.c.v.) in wild-type mice increased the number of CD68+ puncta per microglia (Fig 3L and M) and the percentage of CD68+ microglia (Fig 3N). Thus, EP4 receptor activation stimulates microglial phagocytosis *in vivo* and *in vitro*, providing a basis for the homeostatic (non-phagocytic) microglial phenotype observed with EP4 deficiency.

### Microglial EP4 deficiency preserved POMC neuron projections to the PVN during HFD feeding

Previous studies have demonstrated that HFD feeding triggers alterations to POMC neurons via cell-cell contact with activated microglia and disruption of downstream melanocortin signaling [16]. Given the more quiescent profile of EP4-deficient microglia, we hypothesized that MG-EP4 KO mice maintain more intact POMC neuronal cytoarchitecture during HFD feeding. First, we determined that the overall number of POMC neurons (Fig. 4A and B) and microglia (Fig. 4C) remained unchanged between WT and MG-EP4 KO mice. However, the apposition of POMC neurons and microglial cell bodies was significantly reduced in the MG-EP4 KO group compared to the WT. Specifically, there were fewer points of contact between microglia and POMC neuron soma (marked by β-endorphin immunoreactivity) (Fig. 4D) and POMC processes in the MG-EP4 KO group (Fig. 4E). POMC neurons project to the melanocortin receptor 3/4-expressing neurons in the paraventricular nucleus (PVN), a hypothalamic region critical in food intake control and substrate utilization [1]. Consistent with the reduced POMC contact and decreased phagocytic profile of EP4-deficient microglia, POMC fiber density in the PVN was markedly higher in HFD-fed MG-EP4 KO mice (Fig. 4F and G). Together, these data support a model in which the absence of EP4 limits HFD-induced activation of microglial phagocytosis, thereby preserving POMC projections to the PVN and reducing susceptibility to DIO.

## Discussion

Microglial activation is both necessary and sufficient to promote weight gain and hyperphagia in response to HFD feeding, [6, 16] but the specific drivers of this response have not been fully identified. Here, we present evidence that expression of the PGE2 receptor EP4 by microglia is critical for susceptibility to DIO. Specifically, microglial EP4 deficiency confers protection from HFD-induced hyperphagia, weight gain, and insulin resistance while being dispensable for physiological body weight regulation. At the cellular level, these metabolic improvements correspond with reduced microglial phagocytic activation by HFD, fewer contacts between microglia and POMC neurons, and greater abundance of POMC projections to the PVN. Together, these data implicate EP4-mediated regulation of microglial phagocytosis as an important modulator of energy homeostasis during DIO via impacts on melanocortin circuit integrity.

A recent study implicated hypothalamic PGE2 in microglial activation and metabolic regulation during DIO [9]. Preventing the increase in ventromedial hypothalamic PGE2 during HFD feeding by targeted knock-down of an enzyme involved in the generation of prostaglandin precursor molecules reversed microgliosis and improved glucose tolerance on HFD [9]. Likewise, global deletion of a key enzyme for PGE2 synthesis, mPGES-1, reduced food intake and weight gain on HFD [18]. These reports implicate PGE2 signaling in obesity pathogenesis, but the mechanisms involved remain elusive. Among the four receptors for PGE2, the EP4 receptor is highly expressed throughout the body and exerts various physiological and pathological roles in inflammatory processes, lipid uptake, and vascular functions [7]. Global deletion of EP4 leads to a lipodystrophic phenotype with reduced food intake and weight gain, suggesting that the function of EP4 at the systemic level is dominated by its role in adipose tissue [13]. EP4 is also the primary PGE2 receptor expressed in macrophages where it has been implicated in atherosclerotic lesions in humans and mice [18, 19]. Finally, microglia express among the highest levels of EP4 in the mouse and human brain [20, 21], but prior studies have largely focused on its role in bacterial challenge and Alzheimer’s Disease [10, 11]. Thus, our study is the first to describe a function for brain EP4 signaling in obesity pathogenesis, representing an important step forward in elucidating the involvement of CNS prostaglandins in metabolism.

Our finding that microglial EP4 deletion limits food intake and body weight gain during HFD feeding adds to a small but growing body of literature implicating microgliosis in obesity pathogenesis. Interestingly, two mouse models involving microglial deletion of *Ikk*β and mitochondrial uncoupling protein (*Ucp2*) showed a similar level of obesity resistance as the MG-EP4 KO mice [6, 16]. While the metabolic resilience of these previous mouse lines was ascribed to reduced hypothalamic inflammation, we provide a novel mechanism for the MG-EP4 KO model based on preservation of POMC projections to the PVN during HFD feeding. Similar alterations to the POMC neuron-PVN neurocircuit have been described in the context of aging, intrauterine HFD exposure, and short-term HFD feeding, suggesting that a common pathway involving structural compromise of the melanocortin system may underlie environmentally-induced metabolic dysfunction [22–24]. While the precise mechanism underlying this vulnerability is unknown, the preserved architecture of the melanocortin neurocircuit during DIO in the MG-EP4 KO mice corresponded with fewer phagocytic contacts between microglia and POMC soma and projections. As discussed in more detail below, microglial phagocytosis and its impact on neurocircuit integrity represent a novel mechanism for future studies in the context of metabolic physiology and pathology.

During HFD feeding, the levels of inflammatory cytokines such as IL1β and TNFα as well as markers of phagocytosis such as the myeloid-specific lysosome-associated glycoprotein CD68 increase together with the onset of gliosis in the arcuate nucleus [3, 4, 6, 25]. In MG-EP4 KO mice, there were no changes in inflammatory signaling in DIO despite previous reports that microglial EP4 signaling helps resolve inflammation following lipopolysaccharide challenge [10]. In contrast, CD68 levels were significantly reduced along with microglia displaying a less activated phenotype, characterized by a homeostatic pattern of more branching of projections and an elongated, less ameboid cell body. Thus, these data support the novel finding that the microglial phagocytic responses are regulated separately from the inflammatory cytokine response in the context of DIO.

A reduced microglial phagocytic state in the absence of EP4 signaling may impact synapse remodeling. As the resident phagocytes of the CNS, microglia are responsible for maintaining synaptic integrity. For example, microglial engulfment of presynaptic inputs from retinal ganglion cells via the complement pathway is critical in synapse pruning and stabilization [27]. Likewise, microglial phagocytosis of apoptotic neurons via receptor tyrosine kinases is key in debris clearance during development and injury [28]. Microglia also phagocytose dendritic spines, whole neurons, extracellular matrix components, and myelin debris, all of which may profoundly alter neuronal circuitry and physiological outcomes [26–29]. In the context of pathology, aberrant microglial phagocytosis is associated with disruption of neurocircuits or impairment of neuronal function such as in autism and Alzheimer’s Disease [15, 30]. In fact, deficiency of microglial EP4 disrupts amyloid phagocytosis leading to increased plaque accumulation and plaque density in a mouse model of Alzheimer’s disease [11]. Our current study demonstrates that EP4 deficiency reduces the phagocytic state of microglia, a finding associated with preservation of the melanocortin neurocircuits. Follow-up studies will be critical to delineate the impact of phagocytosis on hypothalamic circuit remodeling during HFD feeding and to identify specific substrates involved for targeting experiments.

In contrast with our study, a previous report using a microglia-specific lipoprotein lipase (Lpl) deletion model found reduced CD68 expression associated with increased body weight gain on a high fat, high sucrose diet [33]. One potential explanation for the discrepancy may be a different substrate of phagocytosis in the context of LPL vs. EP4. Indeed, microglial *Lpl* deletion on HFD was associated with fewer POMC neurons [33], while microglial EP4 deletion in the current study did not alter the total number of POMC neurons (nor did loss of UCP2 [17]). Instead, microglial EP4 deletion resulted in fewer microglial appositions with POMC cell soma and processes, the direct converse of a previous study (from the same authors as the *Lpl* study) showing that increased microglia-POMC neuron contacts are associated with HFD-induced weight gain [16]. The association of these reduced cell-cell contacts with a more intact melanocortin neurocircuit suggests a mechanism linking microglial phagocytosis to the maintenance of POMC neuron function during HFD feeding. Indeed, greater melanocortin fiber density in the PVN has been associated with intact sickness-associated anorexia responses in adult mice [34], while depletion of these projections in mice lacking protein kinase Cλ in POMC neurons results in profound obesity in the context of HFD exposure [35]. Given that POMC neurons comprise a heterogeneous population with diverse functional properties [36], future studies will need to further characterize neuron-microglia appositions in specific POMC (and other neuronal) sub-populations and determine their role in DIO pathogenesis.

## Conclusion

In conclusion, microglial EP4 deletion conferred protection from DIO and obesity-associated insulin resistance. EP4 signaling was associated with a phagocytic response to HFD, and abrogation of this pathway suppressed the microglial phagocytic state yielding fewer microglia-POMC neuron contacts and preserved POMC projections in the PVN. Together, these data implicate PGE2 signaling via EP4 as an important regulator of microglia-POMC neuron interactions during HFD feeding that drives susceptibility to metabolic dysfunction, and provide a basis for investigating microglial phagocytosis as a novel target for obesity therapeutics.

## Acknowledgements

This work was supported by the NIH (R01 DK119754) awarded to JPT and American Heart Association Postdoctoral Fellowship (20POST35210291) awarded to AN. We would like to thank the Nutrition Obesity Research Center (NORC, DK035816) for their assistance with metabolic phenotyping and the Diabetes Research Center (DRC, DK017047) at the University of Washington. The authors declare no conflict of interest.

## Figure Legends

**Supplementary Figure 1: Validation of microglial EP4 deletion and body weight differences on chow.**

(A) Prostaglandin PGE2 receptor EP4 expression (*Ptger4*) in whole hypothalamus and isolated hypothalamic microglia following HFD feeding for a week (n=5-6). (B) Validation of *Ptger4* knockdown in FACS-sorted microglia from WT and MG-EP4 KO mice (n=4). (C) EYFP expression in the CD11b+ and CD11b-cells from the WT and MG-EP4 KO mouse brains. (D) Body weight measurements of WT and MG-EP4 KO mice on chow (n=4-6). Values represent mean ±SEM. *p < .05, Student’s *t test.*

## References

[1] G. S. H. Yeo et al., “The melanocortin pathway and energy homeostasis: From discovery to obesity therapy,” Mol Metab, vol. 48, 2021, doi: 10.1016/j.molmet.2021.101206.

[2] T. L. Horvath et al., “Synaptic input organization of the melanocortin system predicts diet-induced hypothalamic reactive gliosis and obesity,” Proc Natl Acad Sci U S A, vol. 107, no. 33, pp. 14875–14880, 2010, doi: 10.1073/pnas.1004282107.

[3] S. M. Thaler JP, Yi CX, Schur EA, Guyenet SJ, Hwang BH, Dietrich MO, Zhao X, Sarruf DA, Izgur V, Maravilla KR, Nguyen HT, Fischer JD, Matsen ME, Wisse BE, Morton GJ, Horvath TL, Baskin DG, Tschöp MH, “Obesity is associated with hypothalamic injury in rodents and humans.,” J Clin Invest, vol. 122, no. 1, pp. 153–62, 2012, doi: 10.1172/JCI59660.

[4] M. Valdearcos et al., “Microglial Inflammatory Signaling Orchestrates the Hypothalamic Immune Response to Dietary Excess and Mediates Obesity Susceptibility,” Cell Metab, vol. 26, no. 1, pp. 185–197.e3, 2017, doi: 10.1016/j.cmet.2017.05.015.

[5] E. C. Cope et al., “Microglia Play an Active Role in Obesity-Associated Cognitive Decline.,” J Neurosci, vol. 38, no. 41, pp. 8889–8904, Oct. 2018, doi: 10.1523/JNEUROSCI.0789-18.2018.

[6] X.-L. Wang et al., “Microglia-specific knock-down of Bmal1 improves memory and protects mice from high fat diet-induced obesity.,” Mol Psychiatry, May 2021, doi: 10.1038/s41380-021-01169-z.

[7] E. M. Smyth, T. Grosser, M. Wang, Y. Yu, and G. A. FitzGerald, “Prostanoids in health and disease.,” J Lipid Res, vol. 50 Suppl, no. Suppl, pp. S423–8, Apr. 2009, doi: 10.1194/jlr.R800094-JLR200.

[8] A. N. Hata and R. M. Breyer, “Pharmacology and signaling of prostaglandin receptors: multiple roles in inflammation and immune modulation.,” Pharmacol Ther, vol. 103, no. 2, pp. 147–166, Aug. 2004, doi: 10.1016/j.pharmthera.2004.06.003.

[9] M.-L. Lee et al., “Prostaglandin in the ventromedial hypothalamus regulates peripheral glucose metabolism.,” Nat Commun, vol. 12, no. 1, p. 2330, Apr. 2021, doi: 10.1038/s41467-021-22431-6.

[10] J. Shi, J. Johansson, N. S. Woodling, Q. Wang, T. J. Montine, and K. Andreasson, “The prostaglandin E2 E-prostanoid 4 receptor exerts anti-inflammatory effects in brain innate immunity.,” J Immunol, vol. 184, no. 12, pp. 7207–7218, Jun. 2010, doi: 10.4049/jimmunol.0903487.

[11] N. S. Woodling et al., “Suppression of Alzheimer-associated inflammation by microglial prostaglandin-E2 EP4 receptor signaling.,” J Neurosci, vol. 34, no. 17, pp. 5882–5894, Apr. 2014, doi: 10.1523/JNEUROSCI.0410-14.2014.

[12] Y. Esaki et al., “Dual roles of PGE2-EP4 signaling in mouse experimental autoimmune encephalomyelitis.,” Proc Natl Acad Sci U S A, vol. 107, no. 27, pp. 12233–12238, Jul. 2010, doi: 10.1073/pnas.0915112107.

[13] Y. Cai et al., “Mice lacking prostaglandin E receptor subtype 4 manifest disrupted lipid metabolism attributable to impaired triglyceride clearance.,” FASEB J Off Publ Fed Am Soc Exp Biol, vol. 29, no. 12, pp. 4924–4936, Dec. 2015, doi: 10.1096/fj.15-274597.

[14] C. N. Parkhurst et al., “Microglia promote learning-dependent synapse formation through brain-derived neurotrophic factor.,” Cell, vol. 155, no. 7, pp. 1596–1609, Dec. 2013, doi: 10.1016/j.cell.2013.11.030.

[15] S. Hong et al., “Complement and microglia mediate early synapse loss in Alzheimer mouse models,” Science (80-), vol. 352, no. 6286, pp. 712–716, 2016, doi: 10.1126/science.aad8373.

[16] C.-X. Yi et al., “TNFα drives mitochondrial stress in POMC neurons in obesity.,” Nat Commun, vol. 8, p. 15143, May 2017, doi: 10.1038/ncomms15143.

[17] J. D. Kim, N. A. Yoon, S. Jin, and S. Diano, “Microglial UCP2 Mediates Inflammation and Obesity Induced by High-Fat Feeding,” Cell Metab, vol. 30, no. 5, pp. 952–962.e5, 2019, doi: 10.1016/j.cmet.2019.08.010.

[18] C. Pierre et al., “Invalidation of Microsomal Prostaglandin E Synthase-1 (mPGES-1) Reduces Diet-Induced Low-Grade Inflammation and Adiposity.,” Front Physiol, vol. 9, p. 1358, 2018, doi: 10.3389/fphys.2018.01358.

[19] K. Takayama, G. García-Cardena, G. K. Sukhova, J. Comander, M. A. J. Gimbrone, and P. Libby, “Prostaglandin E2 suppresses chemokine production in human macrophages through the EP4 receptor.,” J Biol Chem, vol. 277, no. 46, pp. 44147–44154, Nov. 2002, doi: 10.1074/jbc.M204810200.

[20] V. R. Babaev et al., “Macrophage EP4 deficiency increases apoptosis and suppresses early atherosclerosis.,” Cell Metab, vol. 8, no. 6, pp. 492–501, Dec. 2008, doi: 10.1016/j.cmet.2008.09.005.

[21] Y. Zhang et al., “An RNA-sequencing transcriptome and splicing database of glia, neurons, and vascular cells of the cerebral cortex.,” J Neurosci, vol. 34, no. 36, pp. 11929–11947, Sep. 2014, doi: 10.1523/JNEUROSCI.1860-14.2014.

[22] Y. Zhang et al., “Purification and Characterization of Progenitor and Mature Human Astrocytes Reveals Transcriptional and Functional Differences with Mouse.,” Neuron, vol. 89, no. 1, pp. 37–53, Jan. 2016, doi: 10.1016/j.neuron.2015.11.013.

[23] S.-B. Yang, A.-C. Tien, G. Boddupalli, A. W. Xu, Y. N. Jan, and L. Y. Jan, “Rapamycin ameliorates age-dependent obesity associated with increased mTOR signaling in hypothalamic POMC neurons.,” Neuron, vol. 75, no. 3, pp. 425–436, Aug. 2012, doi: 10.1016/j.neuron.2012.03.043.

[24] M. C. Vogt et al., “Neonatal insulin action impairs hypothalamic neurocircuit formation in response to maternal high-fat feeding.,” Cell, vol. 156, no. 3, pp. 495–509, Jan. 2014, doi: 10.1016/j.cell.2014.01.008.

[25] M. Schneeberger et al., “Reduced α-MSH Underlies Hypothalamic ER-Stress-Induced Hepatic Gluconeogenesis.,” Cell Rep, vol. 12, no. 3, pp. 361–370, Jul. 2015, doi: 10.1016/j.celrep.2015.06.041.

[26] S. Lee, Y. Wu, X. Q. Shi, and J. Zhang, “Characteristics of spinal microglia in aged and obese mice: potential contributions to impaired sensory behavior.,” Immun Ageing, vol. 12, p. 22, 2015, doi: 10.1186/s12979-015-0049-5.

[27] B. Stevens et al., “The classical complement cascade mediates CNS synapse elimination.,” Cell, vol. 131, no. 6, pp. 1164–1178, Dec. 2007, doi: 10.1016/j.cell.2007.10.036.

[28] L. Fourgeaud et al., “TAM receptors regulate multiple features of microglial physiology.,” Nature, vol. 532, no. 7598, pp. 240–244, Apr. 2016, doi: 10.1038/nature17630.

[29] L. Weinhard et al., “Microglia remodel synapses by presynaptic trogocytosis and spine head filopodia induction.,” Nat Commun, vol. 9, no. 1, p. 1228, Mar. 2018, doi: 10.1038/s41467-018-03566-5.

[30] P. T. Nguyen et al., “Microglial Remodeling of the Extracellular Matrix Promotes Synapse Plasticity.,” Cell, vol. 182, no. 2, pp. 388–403.e15, Jul. 2020, doi: 10.1016/j.cell.2020.05.050.

[31] K. Shen et al., “Multiple sclerosis risk gene Mertk is required for microglial activation and subsequent remyelination.,” Cell Rep, vol. 34, no. 10, p. 108835, Mar. 2021, doi: 10.1016/j.celrep.2021.108835.

[32] M. W. Salter and B. Stevens, “Microglia emerge as central players in brain disease.,” Nat Med, vol. 23, no. 9, pp. 1018–1027, Sep. 2017, doi: 10.1038/nm.4397.

[33] Y. Gao et al., “Lipoprotein Lipase Maintains Microglial Innate Immunity in Obesity,” Cell Rep, vol. 20, no. 13, pp. 3034–3042, 2017, doi: 10.1016/j.celrep.2017.09.008.

[34] S. Jin, J. G. Kim, J. W. Park, M. Koch, T. L. Horvath, and B. J. Lee, “Hypothalamic TLR2 triggers sickness behavior via a microglia-neuronal axis.,” Sci Rep, vol. 6, p. 29424, Jul. 2016, doi: 10.1038/srep29424.

[35] M. D. Dorfman et al., “Deletion of protein kinase c l in POMC neurons predisposes to diet-induced obesity,” Diabetes, vol. 66, no. 4, pp. 920–934, 2017, doi: 10.2337/db16-0482.

[36] N. Biglari et al., “Functionally distinct POMC-expressing neuron subpopulations in hypothalamus revealed by intersectional targeting.,” Nat Neurosci, vol. 24, no. 7, pp. 913–929, Jul. 2021, doi: 10.1038/s41593-021-00854-0.

