## Supplementary figures and images for "Prostaglandin PGE2 receptor EP4 regulates microglial phagocytosis and increases susceptibility to diet-induced obesity"

### Supplemental Figure 1

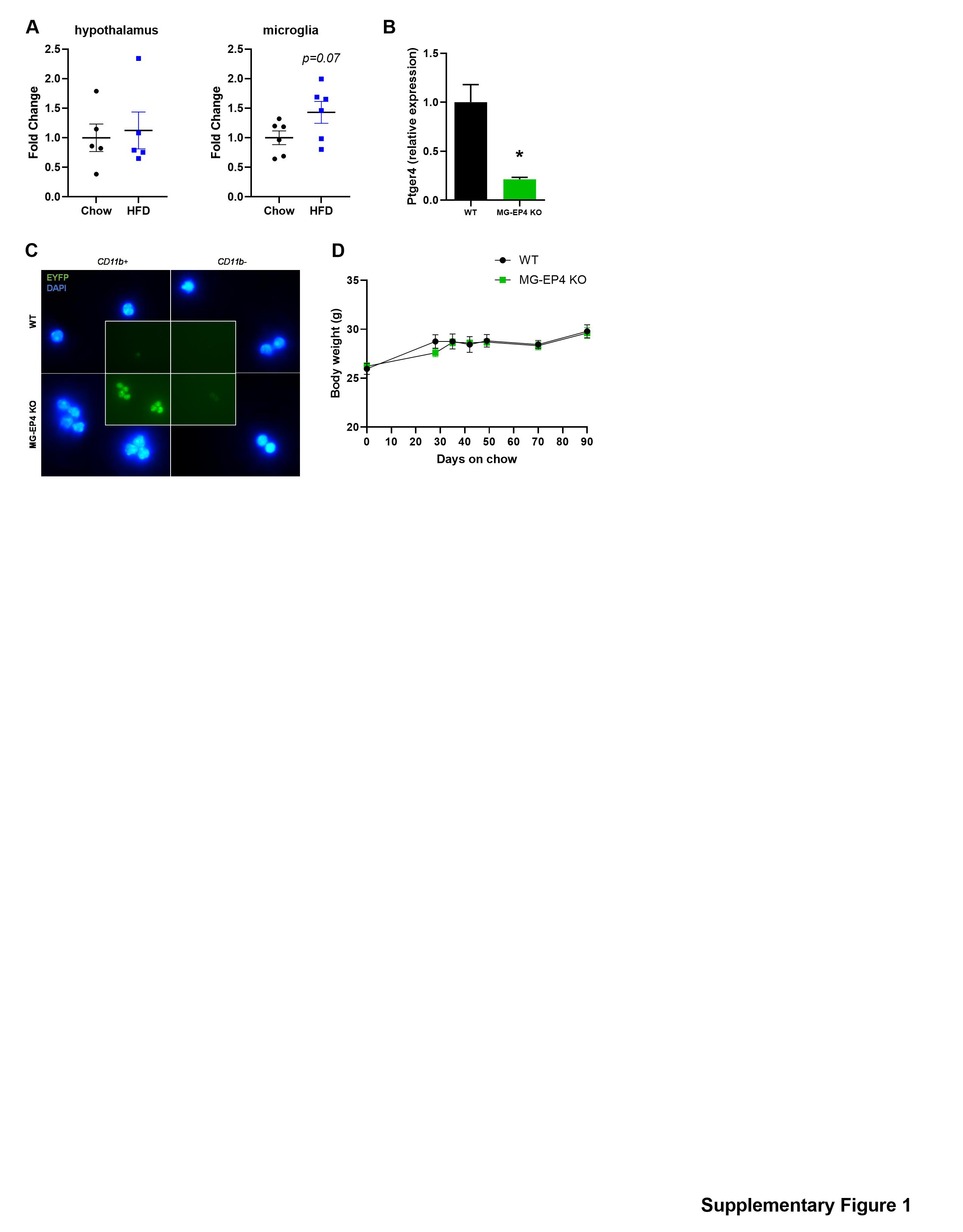
